# *De novo* genome assembly resolving repetitive structures enables genomic analysis of 35 European *Mycoplasma bovis* strains

**DOI:** 10.1101/2023.04.13.536562

**Authors:** Sandra Triebel, Konrad Sachse, Michael Weber, Martin Heller, Celia Diezel, Martin Hölzer, Christiane Schnee, Manja Marz

## Abstract

*Mycoplasma (M*.*) bovis*, the agent of mastitis, pneumonia, and arthritis in cattle, harbors a small genome of approximately 1 Mbp. Combining data from Illumina and Nanopore technologies, we sequenced and assembled the genomes of 35 European strains and isolate DL422_88 from Cuba. While the high proportion of repetitive structures in *M. bovis* genomes represents a particular challenge, implementation of our own pipeline `Mycovista` (available on GitHub www.github.com/sandraTriebel/mycovista) in a hybrid approach enabled contiguous assembly of the genomes and, consequently, improved annotation rates considerably. To put our European strain panel in a global context, we analyzed the new genome sequences together with 175 genome assemblies from public databases. Construction of a phylogenetic tree based on core genes of these 219 strains revealed a clustering pattern according to geographical origin, with European isolates positioned on clades 4 and 5. Genomic data allowing assignment of strains to tissue specificity or certain disease manifestations could not be identified. Seven strains isolated from cattle with systemic circular condition (SCC), still a largely unknown manifestation of *M. bovis* disease, were located on both clades 4 and 5. Pairwise association analysis revealed 108 genomic elements associated with a particular clade of the phylogenetic tree. Further analyzing these hits, 25 genes are functionally annotated and could be linked to a *M. bovis* protein, e.g. various proteases and nucleases, as well as ten variable surface lipoproteins (Vsps) and other surface proteins. These clade-specific genes could serve as useful markers in epidemiological and clinical surveys.

## Introduction

Infections of the bovine pathogen *Mycoplasma (M*.*) bovis* are of considerable economic importance due to their negative impact on animal health and production yields (1). Pneumonia and mastitis are the most prominent clinical manifestations in cattle, but arthritis, genital disorders, and keratoconjunctivitis can also be caused by this agent (2–4). Occasionally, *M. bovis* infections presenting as purulent fibrinous pleuropneumonia with sequestering were observed in adult cattle and had to be differentiated from other *Mycoplasma* diseases, such as Contagious Bovine Pleuropneumonia. Recently, an increase of acute *M. bovis* infections leading to mastitis in medium-sized and large dairy herds was observed in the eastern and northern federal states of Germany. In some cases, a different clinical picture compared to previous outbreaks was noticed since, in addition to mastitis, affected animals showed symptoms of circulatory involvement manifesting as massive edema in the chest, abdomen, and leg area, similar to allergic reactions (see Fig. S1) (5, 6). We will refer to this disease complex as Systemic Circular Condition (SCC).

While many aspects of *M. bovis* pathogenesis are still not fully understood, the agent’s capability of subverting host immune response through surface antigen variation was characterized in several studies (7–10). The central role in this process is played by members of a family of lipoproteins designated variable surface proteins (Vsps), which consist of a conserved N-terminal domain for membrane insertion and lipoprotein processing and a large stretch of repetitive tandem domains comprising up to 80 % of the entire Vsp molecule. As a result of spontaneous and non-coordinated deletions, insertions, and rearrangements in the *vsp* genomic locus, translated Vsps undergo variations in phase (on/off switching), size (varying number of tandem repeats), and/or surface presentation at high frequency (8).

In this context, sequencing and comparative genomics can be important tools to identify genetic features correlating with strain properties and/or disease symptoms (11). High-quality genomes with precise and, if possible, complete annotations are needed to perform meaningful comparative studies. However, despite their small size (∼ 1 Mbp), the assembly of *M. bovis* genomes is challenging due to the high rate of the above-mentioned repetitive structures. In the past, using state-of-the-art tools to assemble Illumina reads resulted in fragmented genomes because repetitive regions are not efficiently covered using short reads. Nevertheless, despite the lower contiguity of such short-read assemblies, the high sequence accuracy of short reads and the associated accurate annotation of open reading frames (ORFs) is essential for further downstream analysis. Currently, 588 assemblies of *M. bovis* strains are available in the NCBI database (Jan 16, 2023), of which 51 assemblies are marked as complete. Nine of them lack GenBank and/or RefSeq annotation entries. The current NCBI reference genome of *M. bovis* (strain 8790, GCF_005061465.1) consists of 17 contigs and is incomplete. Sequencing via nanopores, e.g. the Oxford Nanopore Technologies (ONT) MinION device, has great potential to enable genome assemblies of higher contiguity since longer reads facilitate coverage of complex genomic regions (12, 13). Recent studies have shown that the use of long and short reads is a suitable strategy for assembling bacterial genomes (14).

In the present study, we used a hybrid assembly approach combining the advantages of Illumina and Nanopore sequencing to obtain high-quality genome sequences of 36 *M. bovis* strains. The strain panel contains seven isolates associated with the SCC disease complex in Germany from the last ten years, strains from different geographical regions and from animals with other symptoms, as well as previous isolates. Genome sequence data were processed using our own `Mycovista` pipeline (www.github.com/sandraTriebel/mycovista), which also includes improved annotation, to elucidate epidemiological and phylogenetic relationships.

## Material and Methods

### Strains

The aim was to cover a broad spectrum of *M. bovis* field strains in the study, including isolates from animals with severe symptoms. Therefore, we selected 36 isolates from different geographical regions of Germany, other countries, and different tissue representing various disease manifestations. All German strains isolated in 2014 and later were collected especially for this study by diagnostic laboratories of several federal states and animal health services and incorporated in the strain collection of the Friedrich-Loeffler Institute, Germany. For comparison, the Cuban strain DL422_88 was included. The basic characteristics of the included strains are given in Tab. 1.

### Description of the disease complex

After calving, the Systemic Circular Condition (SCC) affected cows in medium sized and large dairy herds. Animals suffered from painful, swollen joints that partially ruptured, edema mainly in the legs and chest area, shortness of breath, and general nervousness while excreting glassy-serous saliva, usually without any fever. Signs of severe mastitis also typically occurred, with udders hardened to a rubbery consistency, milk becoming creamy and sandy and milk yield dropping sharply. Affected udder quarters either became permanently atrophic or their milk production was fully restored after recovery. In the course of the disease, milk cell counts were significantly elevated. Within a few days, some animals could not stand up and finally died from cardiovascular failure. Treatment with antibiotics had no effect while pain-relieving, anti-inflammatory therapy brought some improvement. Feed consumption was only temporarily reduced. Usually, several animals in the herd fell ill one after another with the same symptoms. For example, on a dairy farm with 450 cows, 60 animals showed the same systemic symptoms, of which 20 died. *M. bovis* was detected in up to 50 % of the diseased cows in a herd. Using commercial ELISA tests, specific antibodies against *M. bovis* were detected in all diseased animals.

### DNA extraction & sequencing

DNA extraction for sequencing with Illumina was done using High Pure PCR Template Preparation Kit from Roche (Mannheim, Germany) according to the manufacturers’ instructions. DNA extraction for Nanopore MinION sequencing was performed using phenol-chloroform extraction as described by Sambrook and Russell (15).

We used four different MinION flow cells (R9.4.1) numbered 1–4 in Tab. S1. Library preparations were done using the 1D genomic DNA by ligation kits (SQK-LSK 108 and SQK-LSK109) and the native barcoding expansion kits (EXP-NBD103, EXP-NBD104, and EXP-NBD114). Briefly, size selection and DNA clean-up were performed using Agencourt AMPure XP beads (Beckman Coulter GmbH, Krefeld, Germany) at a ratio of 1:1 (w:v) before library preparation. Potential nicks in DNA and DNA ends were repaired in a combined step using NEB Next FFPE DNA Repair Mix and NEB Next Ultra II End repair/dA-tailing Module (NewEngland Biolabs, Ipswich, USA) by tripling the incubation time. The ligation of sequencing adapters followed a subsequent second AMPure bead purification onto prepared ends and a third clean-up step with AMPure beads. Additional barcoding and clean-up steps were performed before adapter ligation. Sequencing buffer and loading beads were added to the library. At the start of sequencing, an initial quality check of the flow cells showed 1571, 1379, 1761, and 1236 active pores. Genomic DNA samples used for loading comprised around 50–150 ng per strain (measured by Qubit 4 Fluorometer; ThermoFisher Scientific, Waltham, USA). The sequencing ran for 48 hrs using the MinKNOW software versions 2.2, 18.12.4, 19.05, 19.10. MinION signals are basecalled and demultiplexed using `guppy v6.1.2` model R9.4.1 (only available to ONT customers https://community.nanoporetech.com).

### *De novo* genome assembly

We assembled *M. bovis* genomes using a hybrid approach in our pipeline `Mycovista` (www.github.com/sandraTriebel/mycovista) incorporating both long and short reads. We used `Filtlong v0.2.0` (16) for processing the long reads; `fastp v0.20.0` (17) and `Trimmomatic v0.39` (18) for preprocessing the short reads; `Flye v2.6` (19) for the long read assembly; (i) `Racon v1.3.2` (20), `minimap2 v2.17` (21) and (ii) `medaka v0.11.4` (22) for assembly polishing with long reads, and (iii) `Racon, minimap2` for assembly polishing with short reads; and `Prokka v1.14.5` (23) for genome annotation. A detailed description of our assembly pipeline `Mycovista` can be found in the Results section. We ran `Mycovista` in release version 1.0.

### Genome synteny analysis

We used `Mauve v1.2.0` (24) to detect recombinations in the genome assemblies, such as gene loss, duplication, rearrangement, and horizontal transfer. In order to make our *M. bovis* genome assembly panel comparable regarding genome rearrangements, we designated the first base of the gene *dnaA* to be the first base of the genome as described in the study by Mackiewicz *et al*. (25).

### Pangenome analysis and virulence genes

We used `PPanGGOLiN v1.2.74` (26) for pangenome analysis. `PPanGGOLiN` generates a core gene set based on a Partitioned Pangenome Graph (PPG), which integrates information about protein-coding genes and their genomic neighborhood. The input genomes are annotated by `ppanggolin annotate`. Their specific gene coding table 4 for *Mycoplasma* was set with `--translation_table` 4. Subsequently, clustering was performed using `ppanggolin cluster`, followed by the graph construction and partitioning (`ppanggolin graph, ppanggolin partition`). After generating our 36 assemblies, we compared our data with the study by Yair *et al*. (27), which included 175 assemblies (whole-genome shotgun project PRJNA564939), as well as seven NCBI RefSeq genomes of *M. bovis* (PG45 NC_014760.1, Hubei-1 NC_015725.1, HB0801 NC_018077.1, CQ-W70 NZ_CP005933.1, 08M NZ_CP019639.1, Ningxia-1 NZ_CP023663.1, JF4278 NZ_LT578453.1) (28–32) and one genome provided by the NCBI (NM2012 CP011348.1).

To compare the gene composition of the 35 European strains concerning phylogenetic clustering, we performed statistical association tests (Fisher’s exact test) for the 35 strains using an `R` script (33) available in our `Mycovista` GitHub (www.github.com/sandraTriebel/mycovistatree/master/scripts/gwas.R). More-over, we focused on potential virulence factors (such as Vsps) or genes important for the life cycle (10, 34–36). We listed the presence of these genes based on reference-based annotations. Only genes that were assigned with the correct name for the potential virulence factors were taken into account. Multiple occurrences of a gene are possible due to gene duplication or fragmentation. It is noteworthy that `Prokka` could not annotate *vsp* genes in our assemblies without a reference genome. This may be due to the absence of those sequence annotations in the database or the very diverse structure of *vsp* genes.

**Table 1.**
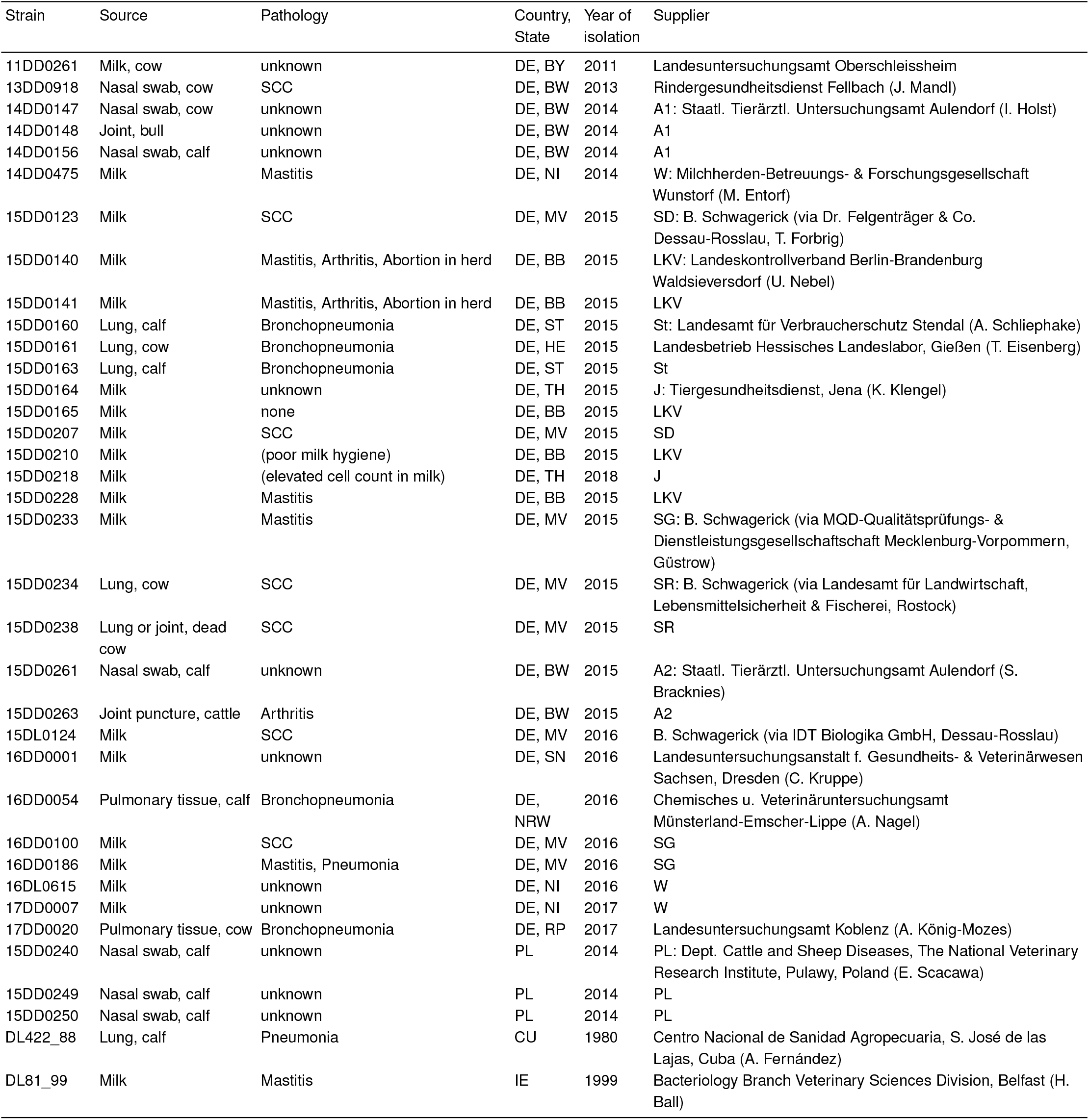
Basic parameters of the *M. bovis* strains included in this study. CU – Cuba; DE – Germany; PL – Poland; IE – Ireland; BB – Brandenburg; BW – Baden-Württemberg; BY – Bavaria; HE – Hesse; SL – Saarland; MV – Mecklenburg Western Pomerania; NI – Lower Saxony; NRW – Northrhine-Westphalia; RP – Rhineland Palatinate; SN – Saxony; ST – Saxony-Anhalt; TH – Thuringia; SCC – Systemic Circulatory Condition;

### Phylogenetic tree reconstruction

Phylogenetic trees were re constructed using IQ-TREE v2.0.3 (37, 38) with a generalized time-reversible (GTR) model based on the core gene set determined in pangenome analysis. For this purpose, we included *M. agalactiae* PG2 (NC_009497.1) as an outgroup (39). The tree reconstruction of our 36 assemblies was done with 1000 bootstrap runs. For comparison among the 219 *M. bovis* strains, we reduced the number of bootstraps to 500. The core genes of each genome were aligned using `ppanggolin msa` with the parameters `--source dna --translation_table 4` and concatenated in one multiple sequence alignment (26).Trees and metadata were visualized using `iTOL v6.6` (40).

## Results

### Hybrid *de novo* assembly pipeline

We developed Mycovista, a pipeline to assemble highly repetitive bacterial genomes such as *M. bovis* in a hybrid or long-read-only sequencing approach. As input, demultiplexed (nanopore) long reads are required. Illumina paired-end short reads can be used in the hybrid assembly mode. `Mycovista` returns the assembled genome, a suitable annotation, and general assembly statistics. The pipeline is automated using the workflow management system `snakemake v7.3.8` (41) in combination with `conda v22.9.0` for reproducible results via stored tool versions in corresponding environment files (42). All code is publicly available on GitHub (www.github.com/sandraTriebel/mycovista).

The pipeline consists of seven steps, see Fig. 1: Before using `Mycovista`, sequencing and basecalling has to be done by the user to provide the input reads. *(0) Sequencing and Basecalling:* The assembly pipeline can be started in two modes: `long` requiring only ONT long reads and `hybrid` where ONT `long` and Illumina paired-end short reads are needed. MinION signals need to be basecalled and demultiplexed using `guppy`. We used `guppy v6.1.2` model R9.4.1. *(1) Raw Read Quality Check:* `FastQC v0.11.8` (43) is incorporated for a first quality check of the raw Illumina data. The ONT raw read quality is visualized with `Nanoplot v1.41.0` (44). *(2) Preprocessing:* Only reads longer than 1,000 nt are considered for the assembly step (filtered via `Filtlong v0.2.0` (16)), as *M. bovis* genomes are known to contain repeats. Illumina raw reads are preprocessed with fastp v0.20.0 (17) for adapter clipping and `Trimmomatic v0.39` (18) (sliding window size 4, Phred score quality cut-off of 28, minimum read length 20) to perform quality trimming. However, using `Trimmomatic` alone is not sufficient to remove all adapter sequences, and thus `fastp` is included. *(3) Preprocessed Read Quality Check:* `FastQC` and `Nanoplot` are again applied to examine the quality of preprocessed reads and to make a comparison to step *(2)* possible. *(4) Assembly:* The filtered long reads are assembled with `Flye v2.6` (19). Optional Illumina reads are not used for assembly but for polishing. *(5) Polishing:* Post-processing of the long-read assemblies consisted of three steps: (i) four polishing runs with `Racon v1.3.2` (20) in combination with `minimap2 v2.17` (21) using long reads followed by (ii) one long read-based polishing with `medaka v0.11.4` (22) and finally (iii) four polishing runs with `Racon` and `minimap2` using short reads. *(6) Assembly Quality Check:* Finally, the quality of the so-produced hybrid assemblies is validated by `QUAST v5.0.2` (45). *(7) Annotation:* All assemblies are annotated using `Prokka v1.14.5` (23) with the parameter `--gcode` 4 to use the required codon table matching characteristics of *M. bovis*.

**Figure 1.**
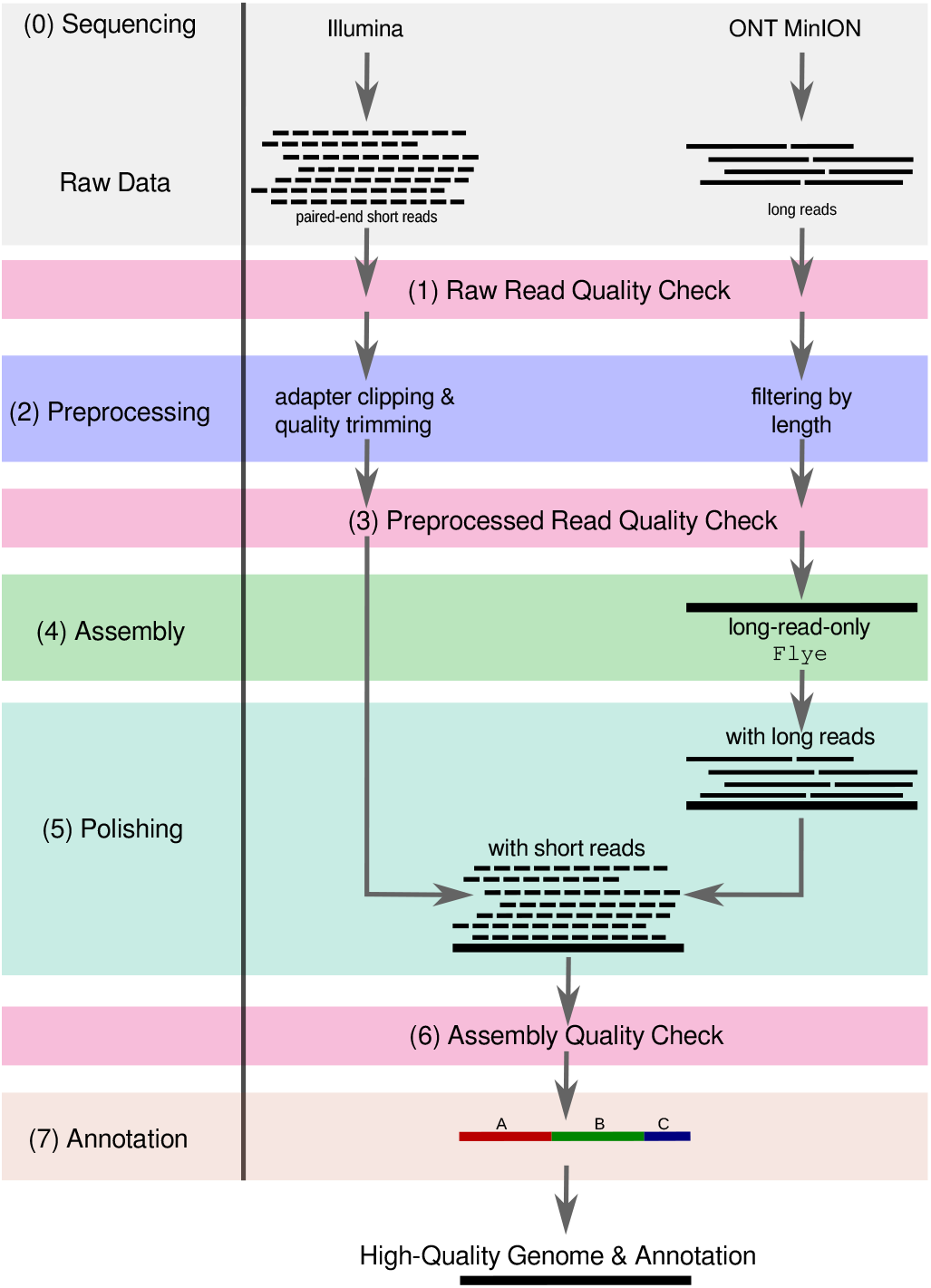
`Mycovista` – a *de novo* assembly pipeline. Mycovista can be used in two assembly modes: long and hybrid, requiring only long reads or additionally paired-end short reads, respectively. *(0)* Basecalled long-read sequencing data (e.g. from ONT) is required. Illumina paired-end short reads can be used as additional input. *(1)* A raw read quality check (`FastQC` and `Nanoplot`) is followed by *(2)* preprocessing of the input reads (`Filtlong, fastp, Trimmomatic`) which are then *(3)* checked regarding their quality. *(4)* Then, a long-read-only assembly is generated by `Flye`.*(5)* Afterwards, the contigs are polished with the preprocessed long reads in several steps (`Racon, minimap2, medaka`). The assembly can be further postprocessed with short reads. Finally, the *(6)* quality of the final assembly is assessed (`QUAST`) followed by a *(7)* gene annotation (`Prokka`).

### Hybrid assembly approach generates contiguous and accurate genomes

Illumina sequencing produced an average of 9,359,418 paired-end reads with a length of 125–150 nt, whereas ONT sequencing produced 317,908 reads on average with a length of about 3,726.8 nt (longest read: 461,771 nt), see Tab. S1 Illumina & ONT Information.

Based on our hybrid approach, `Mycovista` achieved high genome contiguity: All 36 hybrid assemblies are comprised of one to maximal three contigs (with the notable exception of 16 contigs in 17DD0020) and have an average N50 of 1,050,036. This is a remarkable improvement given the challenges posed by multiple sequence repeats in the genome (46). The average genome size of the assemblies in this study was 1.063 Mbp, with the largest genome size being 1.166 Mbp (15DD0218) and the smallest being 0.952 Mbp (15DD0240). Out of 36 *M. bovis* strains, 23 were assembled into single contigs representing the full chromosome. No contigs of plasmid-origin were found by `PlasmidFinder` (47), which is in line with other studies of *M. bovis* (48). The GC content per assembly ranged from 29.11 % to 29.45 % which is in accordance with previously published *M. bovis* genomes.

The number of annotated protein-coding genes in our European strain panel varied between 797 and 962, see Tab. S1, which is in accordance with the expected amount of CDS in *M. bovis* genomes. We observed varying CDS numbers in strains located at different clades of the phylogenetic tree. In the following, we refer to the assemblies according to their association with a particular clade, see Fig. 3, consistent with the literature (27). On average, clade 4 shows 50 more CDS than clades 5 and 6 (clade 4: 910, clade 5: 860, clade 6: 856). Almost every genome consists of six rRNA operons (16S, 5S, and 23S rRNA). The number of tRNAs is 34, whereas the annotation displays two more tRNAs with codons for Arg (CCT, TCT) in 15DD0240. Strain 15DD0263 shows four extra tRNAs with codons for Thr (GGT, CGT), Lys (CTT), and Trp (TCA), respectively. Once per *Mycoplasma* genome, a tmRNA is present, which is representative of other housekeeping ncRNAs illustrating the general quality of the assemblies. Basic statistics about the ONT sequencing, assembly, and annotation results are given in Tab. S1.

### Comparison with global strain panel shows clustering according to geographical origin

To put our assemblies in a global context, we compared the 36 genomes with 175 assemblies from the study by Yair *et al*. (27) and eight complete assemblies of *M. bovis* available from the NCBI database (PG45, Hubei-1, HB0801, CQ-W70, NM2012, 08M, Ningxia-1, JF4278). The phylogenetic tree (see Fig. 2) reconstructed from the core gene set (*M. agalactiae* PG2 included as outgroup) indicates clustering according to geographical origin and is in agreement with the findings of Yair *et al*. With the exception of the Cuban strain DL422_88 (marked as clade 0), all of our *de novo* assembled genomes are located in the European cluster (clades 4, 5, and 6), which is consistent with their origin.

**Figure 2.**
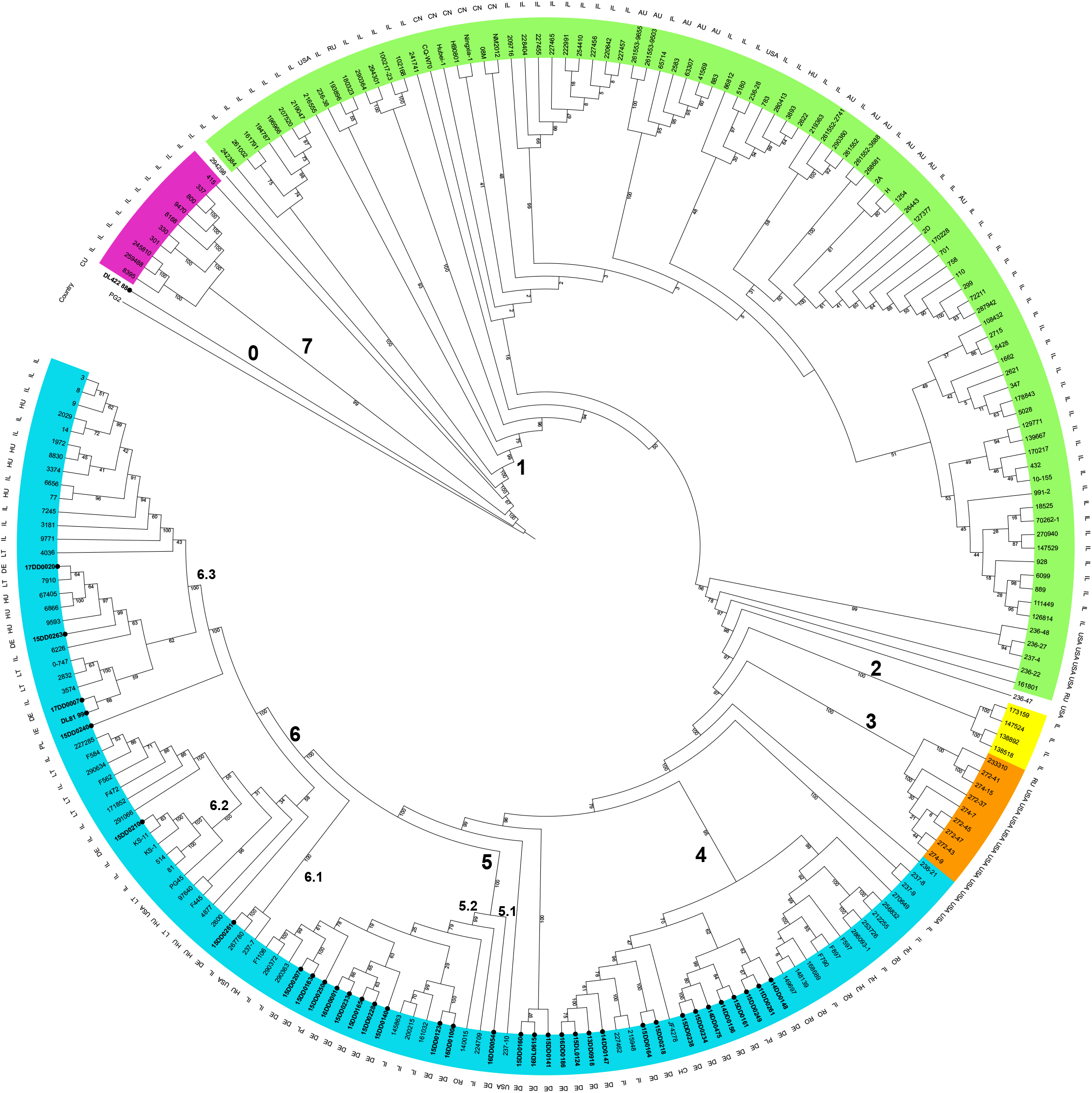
Phylogeny of 219 *M. bovis* isolates based on the core gene set. We combined our *de novo* assembled strain panel with 175 assemblies deposited at the whole genome shotgun project PRJNA564939 (27) and eight complete genomes of *M. bovis* available at the NCBI database. *M. agalactiae* PG2 was included as an outgroup. The clustering of genomes reflects the geographical origin of strains, with 35 of our own strain panel situated in clades 4, 5, and 6 of the European cluster (cyan), and Cuban strain DL422_88 standing outside. Clade numbers and colors are the same as in the SNP-based phylogenetic tree of the paper by Yair *et al*. (27).

For a more detailed analysis of our *de novo* assembled strain panel, we reconstructed a phylogenetic tree based on the genomes (Fig. 3). The tree reveals the same clustering pattern as observed in the global phylogenetic tree in Fig. 2. Therefore, we will use the same clade designations 0, 4, 5, and 6 when referring to members of our strain panel.

**Figure 3.**
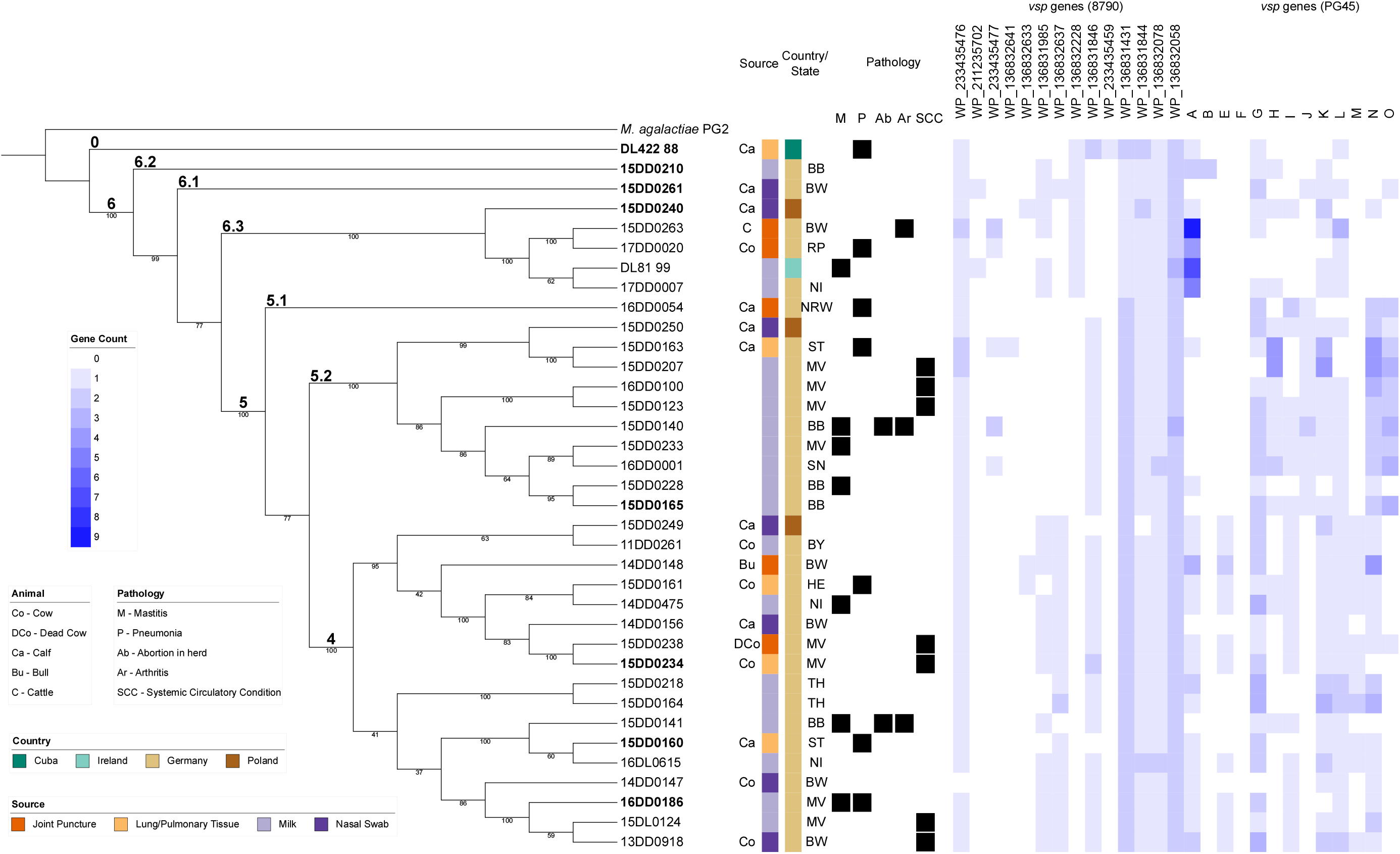
Phylogenetic tree of our *de novo* assembled strain panel. The tree can be divided into three major clusters (4, 5, and 6) and one outlier (Cuban strain DL422_88). We included *M. agalactiae* PG2 as an outgroup. Biological information such as geographical origin (country, state), source (animal, tissue), pathology, and the presence/absence of *vsp* genes are shown. Strain designations in boldface denote representative sequences of the clusters used for genome synteny analysis shown in Fig. 4. In the right-hand part, *vsp* genes in each strain annotated according to NCBI RefSeq strains 8790 and PG45 are depicted. Clade numbers are labeled according to the study by Yair *et al*. (27). Bio Information – Biological Information: Central columns show source of isolation (animal, tissue), geographical origin (country, state), and pathology (if available); Co – Cow; DCo – Dead Cow; Ca – Calf; Bu – Bull C – Cattle; BB – Brandenburg; BW – Baden Wurttemberg; BY – Bavaria; HE – Hesse; SL – Saarland; MV – Mecklenburg Western Pomerania; NI – Lower Saxony; NRW – Northrhine-Westphalia; RP – Rhineland Palatinate; SN – Saxony; ST – Saxony-Anhalt; TH – Thuringia; M – Mastitis; P – Pneumonia/Bronchopneumonia; Ab – Abortions; Ar – Arthritis; SCC – Systemic Circulatory Condition;

We observed no enrichment of a single clade for isolation source or geographical parameters. Concerning disease manifestations, it is noteworthy that the seven strains associated with SCC were located in two clades: 5.2 and 4. As shown in the right-hand part of Fig. 3, the *vsp* gene content varies considerably among the *M. bovis* strains. We used two different sets of annotated variable surface proteins based on strains 8790 (GCF_005061465.1) and PG45 (NC_014760.1), respectively, to search for homologs in our strains. Analysis of the reference-based annotations revealed three general patterns characterizing the occurrence of individual *vsp* genes in the European strain panel: *i)* abundant *vsps* (occurring in all or almost all strains), *ii)* rare *vsps* (occurring in a few strains only or absent altogether), as well as *iii)* those having a preference for one or two clades. Five of the strain 8790-based *vsp* genes were classified as abundant (WP_233435476, WP_136831431, WP_136831844, WP_136832078, WP_136832058), four as rare (WP_211235702, WP_136832641, WP_136832633, WP_233435459), and the following genes showed a clade preference: WP_233435477 (for clades 5 and 6), WP_136831 985 (4 and 6), WP_136832637 (4), WP_136832228 (6), and WP_136831846 (5). When looking for *vsp* homologs of strain PG45, *vsps* G, K, L and N were present in almost all strains. Members of subclade 6.3 were found to harbor several *vspA* gene copies in their genome. The *vspB* gene was identified only in one strain, while *vspF* was absent in all 36 strains. Among *vsps* with clade preference, *vspE* and *vspM* genes seemed to be confined to clade 4, whereas *vspJ* and *vspO* were only encountered in clade 5.

### Comparison of genome synteny among the European strain panel

Analysis of the global organization of the 36 *de novo* assembled genomes revealed rearrangements when comparing clusters or subclusters. Notably, there are characteristic features that are consistent within the genomes of a cluster, which also indicates that they are not the result of assembly errors. To illustrate clade-specific changes in genome synteny, we selected eight strains representing the major clades for calculating an alignment using `Mauve`, see Fig 4.

**Figure 4.**
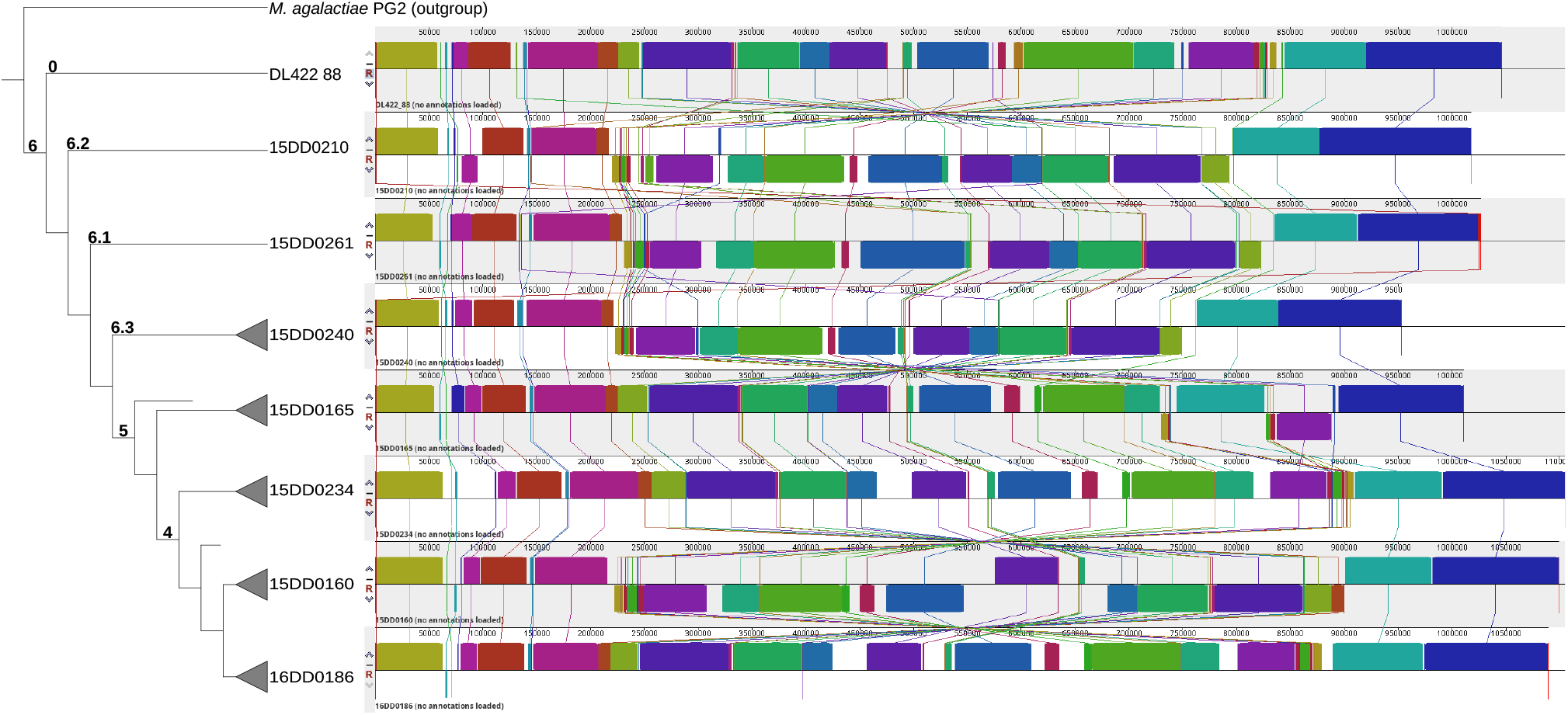
Multiple genome alignment of eight strains representing the major clades using Mauve. Each genome was linearized and normalized, with homologous segments (locally collinear blocks) shown as colored rectangles. Inverted regions are set below those that match the neighboring genome in the forward orientation. Lines collate aligned segments between genomes. The alignment of all 36 assemblies done in this study is shown in Fig. S2.

While the first three blocks 5’ and the last two blocks 3’ are highly conserved among all strains in terms of synteny, the central genomic region, which contains more than 10 locally collinear blocks, displays the highest divergence. As expected, the Cuban strain DL422_88 is distinct from all other European isolates and has a markedly different synteny compared to the rest of the strain panel. A major characteristic of clade 6 strains (15DD0210, 15DD0261, 15DD0240) is the inverted central part of the genome, in which the blocks appear in the reverse complement orientation relative to the other genomes. This region displays the highest degree of divergence among clade members in comparison to clades 4 and 5 (Fig. S2). Strain 15DD0160 (and two other strains from clade 4, see fig. S2) also appears to have an inverted central block, similar to the strains of clade 6. In contrast, the remaining representatives of clades 4 (15DD0234 and 16DD0186) and 5 (15DD0165) share the central genomic blocks arranged in the forward orientation, see Fig. 4. There is also variation among individual strains within the clade (see Fig. S2). The genomes of clade 4 are 50,000 to 100,000 bp larger than those of clades 5 and 6, which explains the higher number of CDSs in clade 4 strains. These genomes contain a number of segments depicted as blank spaces that have no homologs in the other strains (Fig. 4 and S2).

### Pangenome analysis reveals cluster-associated genes including virulence factors

We identified a pangenome of our 36 assemblies consisting of 1,143 gene families, composed of 598 core genes (i.e., genes present in all 36 genomes), 105 shell genes present in ∼55 % genomes, and 351 cloud genes present at low frequency (∼8 %). Gene association analysis revealed 108 to be significantly associated with at least one cluster (4, 5, 6), i.e. are either significantly enriched or depleted in the strains of the analyzed cluster (see Fig. S3). Of those genes, 25 could be linked to a gene product or at least a class of products. These clade-specific genes include various enzymes, e.g. proteases and nucleases, as well as ten variable surface lipoproteins and other surface proteins (see Fig. 5). In particular, some *vsp* genes show correlations to phylogenetic clusters. In Fig. 3, we can observe *vsp WP_136832228* to be only present in clade 6 and Cuban strain DL422_88 (clade 0). The genes *vspJ* and *vspO* are only present in clade 5. Genes *vsp WP_136832637* and *vspE* are associated with clade 4.

**Figure 5.**
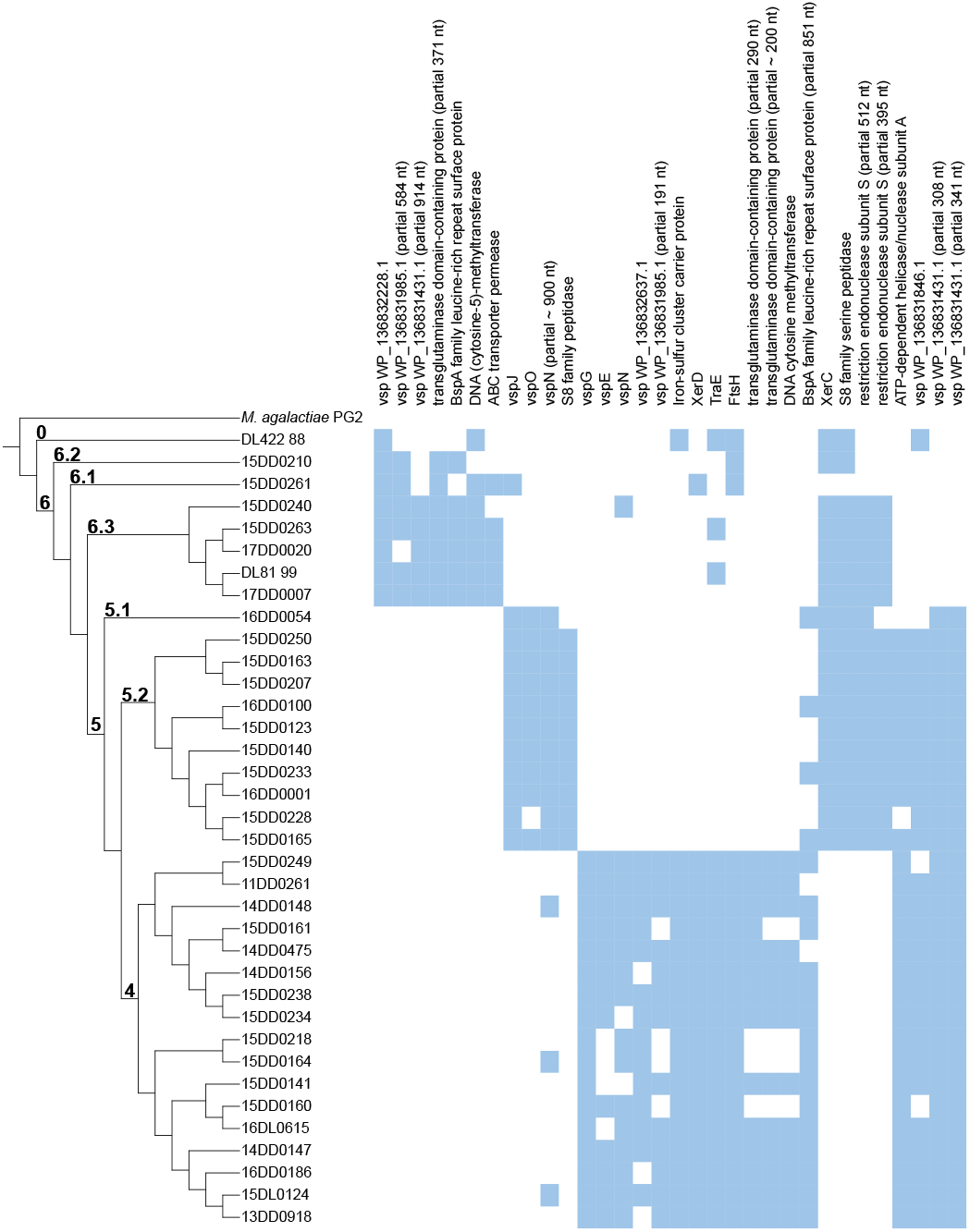
Gene association analysis (Fisher’s exact test) of our European assembly panel in correlation to phylogeny. We detected 108 genes to be significantly associated with at least one cluster. Of these genes, 25 are linked to a gene product or at least a class of products, including various enzymes, e.g. proteases and nucleases, as well as ten variable surface lipoproteins and other surface proteins. Some genes are listed more than once when truncated versions are also associated with a cluster.

As shown in Fig. 5, genes encoding iron-sulfur cluster carrier protein, tyrosine recombinase XerD, conjugal transfer protein TraE, ATP-dependent zinc metalloprotease FtsH, transglutaminase domain-containing protein, and DNA cytosine methyltransferase seem to be only annotated in genomes of clade 4. Gene *xerC* coding for tyrosine recombinase XerC is annotated three to four times in clades 5 and 6, but only two times in clade 4. We observed these differences in the annotation files as well as in the gene association analysis in which one annotation of *xerC* was detected as absent in clade 4. The gene *vsp WP_136831846* is absent in clade 6. Genes *vspN* were annotated multiple times in some genomes. One of them occurred only in clade 5 strains, and another one was only present in clade 4. We observed similar results for *vsp WP_136831985*, see Fig. 5, which was never annotated in group 5 genomes, but found once in each of clades 4 and 6, respectively. The gene *vsp WP_136831431* was annotated twice in clades 4 and 5 and in Cuban strain DL422_88 (clade 0), but only once in clade 6 as revealed by Fisher’s exact test.

## Discussion

### *De novo* assembly

`Mycovista` assembles bacterial genomes using a hybrid approach to tackle the problem of resolving highly repetitive regions. The use of long reads (ONT used here; in theory PacBio too, but not tested) in the initial assembly is beneficial for generating contiguous genomes, and the polishing steps involving short reads improve the sequence accuracy of the genomes. This aligns with state-of-the-art studies and current best practices in assembling high-quality bacterial genomes (14). General assembly statistics and genome annotation are additional steps in the pipeline for further analysis. With our method, we were able to assemble 36 *M. bovis* strains examined in this study into contiguous genomes and to ensure high sequence accuracy through postprocessing of the assemblies, resulting in suitable annotations.

### Phylogenetic clades

Comparison in a global context showed clustering according to geographical origin, with the newly assembled genomes positioned in the European cluster (comprising clades 4, 5, and 6) of the tree. In contrast, the genome of the Cuban strain DL422_88 proved phylogenetically more distant and did not cluster with any other genome.

The geographic clustering, however, is not of high resolution, i.e. not country-specific, which is probably due to the extensive international animal trade throughout Europe. The fact that the European cluster also includes strains from the US (e.g. type strain PG45) could be seen as an indication that phylogenetic processes in *M. bovis* occur at a slow pace.

Analysis of our assemblies also revealed cluster association of several genes, including the *vsps*, whose products are presumed virulence factors. Other correlations between phylogeny and strain properties were not observed. Genomic data supporting tissue specificity of strains or association with specific disease manifestations have not been provided in the literature so far. The identification of a mastitis-dominant lineage of *M. bovis* strains by Yair *et al*. (27) was certainly due to the relatively small size of the cattle population in Israel, where calves were imported from five to seven countries only. Nevertheless, strains associated with mastitis from that study did not cluster on a single clade, which indicates that genomic factors alone are probably insufficient to characterize a strain’s association with a particular clinical manifestation.

### Genome, core genome, pangenome

Genomes of *Mycoplasma* spp. are the smallest among bacteria. With an average length of 1.063 Mbp, our panel of 36 strains has genomes of the expected size, with a core genome of 598 genes and a pangenome of 1,143 (Tab. S1). The average number of annotated protein-coding genes of 880 (Tab. S1) is higher than in other studies, probably due to the hybrid assembly approach, which allowed complete genome assembly in most cases and improved annotation. Kumar *et al*. analyzed a large set of 250 *M. bovis* strains, mainly from the US and Australia, appeared to be phylogenetically more heterogeneous than our European panel. These authors used Illumina sequence data and reported an average CDS number of 770, core and pangenome of 283 and 1,186, respectively (49). The data of our own study comprising 219 strains are comparable: 327 core genes and 1,623 genes in the pangenome (the latter due to annotation, see above).

### Essential vs. dispensable genes

From a formal point of view, the core genome elements are considered indispensable for the organism’s survival (50). From a biological viewpoint, genes encoding replication and translation factors, as well as elements of metabolic pathways, are considered essential because they belong to the fundamental cellular machinery of bacteria. Josi *et al*. (51) identified 352 out of 900 *M. bovis* genes as essential, most of them involved in nucleotide metabolism or biosynthesis of secondary metabolites and amino acids. Using this definition, the clade-specific genes identified in our study (partly given in Fig. 5) would mainly be classified as non-essential.

### *vsp* locus

No precise figures on the varying size of the entire *vsp* locus in *M. bovis* genomes are available. Early data from Lysnyansky *et al*. (7, 8) indicate that the size of all 13 sequences coding for Vsp polypeptide chains in strain PG45 is about 11 kbp, to which the promoter (150 bp each) and signal peptide (75 bp) regions have to be added (total approx. 3 kbp). This means that 1.4 % of the genome would be used to encode Vsp family members. The proportion could be higher in strains equipped with multiple *vsp* gene copies and lower in strains lacking individual *vsp* genes.

Josi *et al*. classified *vsp* genes as non-essential for the organism (51). This may be due to the software used in their study, which could not assign reads from highly repetitive regions to a specific position in the genome, and, therefore, did not consider *vsps*. From the biological perspective, it seems more likely that the *vsp* locus is not dispensable since individual members were suggested to play a role in cytoadhesion and evasion of the host immune response (10, 52). The fact that *M. bovis* strains can swiftly alter their Vsp repertoire and protein chain length in the face of host or environmental challenges could also indicate that the underlying genes are essential for the organism.

When Kumar *et al*. used the sequences of the 13 *vsp* genes of strain PG45 as BLAST queries they found that none of their strains harbored the complete *vsp* gene set (49). While it is known that the number of *vsp* family members can be reduced in certain *M. bovis* strains the high degree of sequence variation in the *vsp* genes may have contributed to the low number of hits in that study.

### SCC as a largely unknown manifestation of *M. bovis* infection

The symptomatology observed in SCC has not been described before and represents an additional clinical picture that might be associated with *M. bovis* infection. The signs of this new condition primarily include severe edema, particularly in the thoracic and abdominal regions, as well as arthritis, even in adult animals (see Fig. S1). The formation of edema indicates involvement of the cardiovascular system. We observed several fatalities in dairy cows with SCC in conjunction with *M. bovis* isolation. Our hypothesis was that *M. bovis* strains isolated from animals presenting with SCC carry genomic traits responsible for higher virulence. However, using our comparative genomic approach involving genome-wide association analysis, such traits could not be identified. Therefore, the question of whether *M. bovis* contributes directly or indirectly to SCC pathogenesis remains open. In addition to SCC, several cases have been reported, in which *M. bovis* was not only implicated in Bovine Respiratory Disease (BRD) in calves but also caused fibrinous bronchopneumonia in adult animals reminiscent of Contagious Bovine Pleuropneumonia (CBPP) (5, 6). This underlines the importance of *M. bovis* as a differential diagnosis for the reportable CBPP, which should be conducted by means of molecular identification tools.

## Acknowledgements

We thank all the colleagues mentioned in Tab. 1, who kindly provided strains for this study. Susann Bahrmann is acknowledged for excellent technical assistance.

## Availability of data and materials

`Mycovista` is available on GitHub https://github.com/sandraTriebel/mycovista. Our *de novo* assembly panel (36 assemblies) is provided in the NCBI BioProject PRJNA954308.

## Authors’ contributions

KS, MHö, and MM designed the study. KS did conventional sequence assemblies at the initial stage. MHö and ST developed the new assembly pipeline. ST conducted the assembly and processing of raw sequence data. MHe and CS provided the strains and submitted them for sequencing. CD did Nanopore sequencing and wetlab work. MW conducte pairwise association analysis. ST, KS, MHö, and MM wrote the manuscript. All authors read and approved the final version of the manuscript.

