## Supplementary Information for "*De novo* genome assembly resolving repetitive structures enables genomic analysis of 35 European *Mycoplasma bovis* strains"

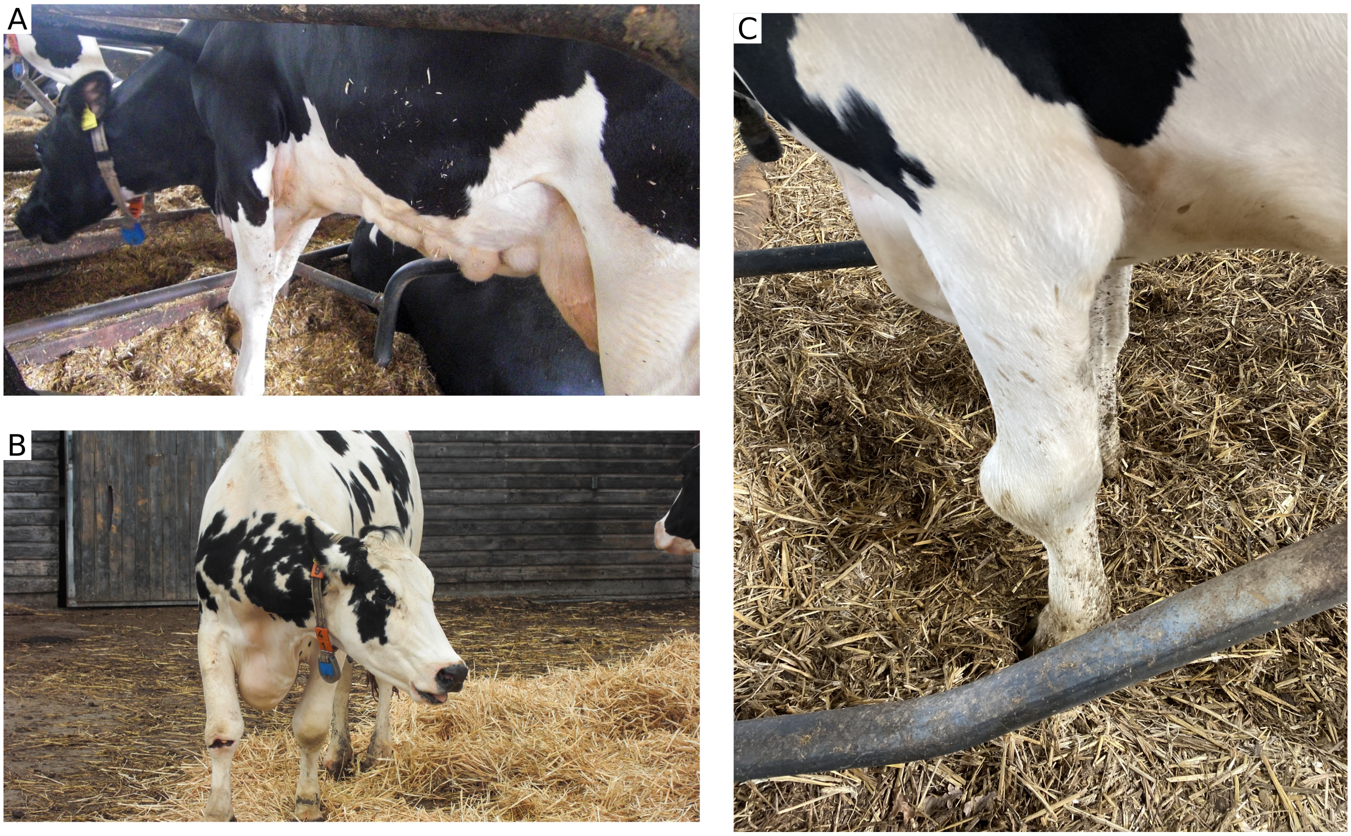

**Figure S1.** *M. bovis* infection causing systemic circular condition (SCC) showing signs of edema and arthritis. (A) Edema in the abdominal area, (B) Severe edema in the chest area, (C) Arthitis of the front leg.

**Table S1.** General assembly statistics of 36 *M. bovis* strains. Bio Information – Biological Information #Reads – number of raw reads; MeanL – mean length of raw reads; F – flow cell number; LR – length of longest read (and its quality) in bp; P – processed; #C – number of contigs after assembly; Total L – length of total assembly in bp; L Contg – length of longest contig in bp; GC – GC content (%); CDS – number of annotated CDS (including unknown proteins); CDS* – number of CDS with annotated gene name (without hypothetical proteins and putative product names); tR – number of tRNAs; rR – number of rRNAs; tm – number of tmRNAs;

| Bio Information | | | | Illumina Information | | ONT Information | | | | | General Assembly Statistics | | | | Annotation Statistics | | | | |
|---|---|---|---|---|---|---|---|---|---|---|---|---|---|---|---|---|---|---|---|
| | | Year | Clade | #Reads | MeanL | F | #Reads | MeanL | N50 | LR (quality) | #C | Total L | L Contg | GC | CDS | CDS* | tR | rR | tm |
| Ref 8790 | | | | | | | | | | | 17 | 905,797 | 254,357 | 29.50 | 742 | | 34 | 4 | 1 |
| Ref PG45 | | | 6.2 | | | | | | | | 1 | 1,003,404 | 1,003,404 | 29.30 | 829 | | 34 | 6 | |
| DL422_88 | Ca | 1988 | 0 | 22,148,828 | 126 | 4 | 249,877 | 7,129.1 | 16,243 | 133,702 (14.2) | 1 | 1,046,374 | 1,046,374 | 29.20 | 871 | 355 | 34 | 6 | 1 |
| 15DD0210 | | 2015 | 6.2 | 11,356,887 | 126 | 3 | 491,868 | 2,429.0 | 4,334 | 109,119 ( 9.4) | 1 | 1,017,740 | 1,017,740 | 29.29 | 853 | 364 | 34 | 6 | 1 |
| 15DD0261 | Ca | 2015 | 6.1 | 9,552,559 | 151 | 3 | 364,896 | 1,884.8 | 4,170 | 102,976 ( 9.2) | 1 | 1,026,147 | 1,026,147 | 29.30 | 856 | 354 | 34 | 6 | 1 |
| 15DD0240 | Ca | 2014 | | 7,709,202 | 126 | 4 | 204,408 | 3,728.3 | 14,068 | 227,226 (11.1) | 1 | 953,284 | 953,284 | 29.37 | 797 | 354 | 36 | 6 | 1 |
| 15DD0263 | C | 2015 | | 12,082,542 | 126 | 4 | 180,976 | 4,174.2 | 12,812 | 180,080 (10.5) | 3 | 1,046,052 | 908,152 | 29.39 | 888 | 371 | 38 | 6 | 1 |
| 17DD0020 | Co | 2017 | 6.3 | 6,893,910 | 151 | 1 | 229,537 | 803.0 | 985 | 41,015 (11.5) | 16 | 990,775 | 330,523 | 29.25 | 855 | 356 | 34 | 4 | 1 |
| DL81_99 | | 1999 | | 8,215,831 | 151 | 4 | 240,108 | 2,722.5 | 5,861 | 122,336 ( 9.3) | 1 | 1,044,487 | 1,044,487 | 29.41 | 888 | 352 | 34 | 6 | 1 |
| 17DD0007 | | 2017 | | 7,165,832 | 151 | 1 | 369,101 | 732.9 | 908 | 42,771 (11.8) | 1 | 991,212 | 991,212 | 29.27 | 837 | 351 | 34 | 4 | 1 |
| 16DD0054 | Ca | 2016 | | 1,716,256 | 126 | 4 | 241,014 | 3,434.5 | 5,438 | 132,662 ( 9.8) | 1 | 996,378 | 996,378 | 29.43 | 825 | 345 | 34 | 6 | 1 |
| 15DD0250 | Ca | 2014 | | 13,235,007 | 126 | 4 | 125,566 | 4,476.8 | 14,729 | 170,021 (13.8) | 1 | 1,027,924 | 1,027,924 | 29.35 | 860 | 354 | 34 | 6 | 1 |
| 15DD0163 | Ca | 2015 | | 8,316,899 | 126 | 3 | 98,914 | 5,206.0 | 12,346 | 125,285 (14.5) | 2 | 1,045,717 | 1,040,623 | 29.36 | 870 | 357 | 35 | 6 | 1 |
| 15DD0207 | | 2015 | | 10,222,109 | 126 | 3 | 252,526 | 4,304.4 | 8,400 | 121,699 ( 9.2) | 1 | 1,053,933 | 1,053,933 | 29.36 | 875 | 361 | 35 | 6 | 1 |
| 16DD0100 | | 2016 | | 8,085,183 | 126 | 4 | 116,974 | 6,963.2 | 18,046 | 210,665 (11.1) | 1 | 1,002,935 | 1,002,935 | 29.34 | 835 | 351 | 34 | 6 | 1 |
| 15DD0123 | | 2015 | 5 | 8,815,543 | 126 | 2 | 257,173 | 4,534.8 | 10,474 | 170,872 (10.4) | 1 | 1,010,945 | 1,010,945 | 29.35 | 841 | 351 | 34 | 6 | 1 |
| 15DD0140 | | 2015 | | 7,206,604 | 126 | 2 | 500,382 | 3,080.2 | 6,389 | 152,850 ( 9.6) | 2 | 1,015,460 | 986,739 | 29.41 | 850 | 350 | 34 | 6 | 1 |
| 15DD0233 | | 2015 | | 5,987,289 | 126 | 3 | 177,766 | 3,550.0 | 7,443 | 100,036 (10.0) | 1 | 1,005,274 | 1,005,274 | 29.36 | 840 | 348 | 34 | 6 | 1 |
| 16DD0001 | | 2016 | | 5,497,038 | 151 | 4 | 172,515 | 4,334.7 | 12,277 | 147,403 ( 9.3) | 1 | 1,011,582 | 1,011,582 | 29.37 | 843 | 351 | 34 | 6 | 1 |
| 15DD0228 | | 2015 | | 7,896,760 | 126 | 3 | 596,225 | 1,696.5 | 3,687 | 161,386 ( 3.7) | 1 | 971,138 | 971,138 | 29.34 | 804 | 349 | 34 | 6 | 1 |
| 15DD0165 | | 2015 | | 7,013,238 | 126 | 3 | 323,361 | 2,961.5 | 5,992 | 133,582 (13.0) | 1 | 1,010,634 | 1,010,634 | 29.39 | 848 | 351 | 34 | 6 | 1 |
| 15DD0249 | Ca | 2014 | | 7,483,705 | 126 | 4 | 179,761 | 6,841.2 | 18,142 | 188,317 (10.2) | 1 | 1,065,918 | 1,065,918 | 29.22 | 890 | 352 | 34 | 6 | 1 |
| 11DD0261 | Co | 2011 | | 9,498,037 | 126 | 2 | 263,161 | 4,678.6 | 8,786 | 140,830 ( 8.6) | 1 | 1,122,166 | 1,122,166 | 29.15 | 932 | 360 | 34 | 6 | 1 |
| 14DD0148 | Bu | 2014 | | 10,052,897 | 126 | 2 | 60,465 | 2,836.0 | 6,760 | 120,138 (14.1) | 2 | 1,095,392 | 1,090,223 | 29.22 | 907 | 353 | 35 | 6 | 1 |
| 15DD0161 | Co | 2015 | | 7,910,036 | 126 | 2 | 472,501 | 3,305.6 | 8,117 | 320,318 ( 5.5) | 1 | 1,091,493 | 1,091,493 | 29.19 | 912 | 352 | 34 | 6 | 1 |
| 14DD0475 | | 2014 | | 7,181,419 | 151 | 2 | 525,440 | 2,133.0 | 3,920 | 173,043 (12.2) | 1 | 1,077,003 | 1,077,003 | 29.28 | 896 | 353 | 34 | 8 | 1 |
| 14DD0156 | Ca | 2014 | | 10,609,103 | 126 | 2 | 909,463 | 3,103.3 | 5,784 | 461,771 ( 7.2) | 4 | 1,073,809 | 576,443 | 29.19 | 889 | 354 | 34 | 6 | 1 |
| 15DD0238 | DCo | 2015 | | 11,278,154 | 126 | 3 | 608,083 | 2,098.8 | 3,761 | 127,525 (14.3) | 1 | 1,091,568 | 1,091,568 | 29.19 | 908 | 352 | 34 | 6 | 1 |
| 15DD0234 | Co | 2015 | | 10,832,930 | 126 | 3 | 215,402 | 3,602.3 | 8,541 | 119,027 (10.3) | 1 | 1,105,287 | 1,105,287 | 29.13 | 919 | 352 | 34 | 6 | 1 |
| 15DD0218 | | 2018 | 4 | 10,443,706 | 126 | 3 | 419,865 | 1,672.5 | 2,671 | 124,420 ( 9.8) | 1 | 1,135,727 | 1,135,727 | 29.16 | 946 | 352 | 34 | 8 | 1 |
| 15DD0164 | | 2015 | | 9,059,621 | 126 | 3 | 952,993 | 1,966.5 | 3,699 | 116,403 (12.1) | 1 | 1,133,513 | 1,133,513 | 29.15 | 952 | 352 | 34 | 6 | 1 |
| 15DD0141 | | 2015 | | 8,812,480 | 126 | 2 | 427,464 | 3,600.9 | 7,303 | 170,001 (14.7) | 1 | 1,068,782 | 1,068,782 | 29.21 | 899 | 353 | 34 | 6 | 1 |
| 15DD0160 | Ca | 2015 | | 10,174,035 | 126 | 2 | 349,154 | 3,713.5 | 7,441 | 156,846 (13.4) | 1 | 1,098,927 | 1,098,927 | 29.20 | 920 | 353 | 34 | 6 | 1 |
| 16DL0615 | | 2016 | | 8,347,668 | 151 | 4 | 233,156 | 3,437.4 | 9,291 | 124,555 (12.2) | 1 | 1,141,547 | 1,141,547 | 29.13 | 962 | 358 | 34 | 6 | 1 |
| 14DD0147 | Co | 2014 | | 10,294,081 | 126 | 3 | 307,197 | 5,333.0 | 13,275 | 315,817 (12.4) | 1 | 1,071,008 | 1,071,008 | 29.22 | 892 | 352 | 34 | 6 | 1 |
| 16DD0186 | | 2016 | | 13,582,089 | 126 | 4 | 193,932 | 6,233.6 | 14,319 | 178,119 (12.2) | 1 | 1,089,251 | 1,089,251 | 29.19 | 912 | 352 | 34 | 6 | 1 |
| 15DL0124 | | 2015 | | 10,749,259 | 151 | 4 | 94,911 | 6,966.1 | 16,334 | 169,771 ( 9.5) | 1 | 1,081,338 | 1,081,338 | 29.19 | 906 | 352 | 34 | 6 | 1 |
| 13DD0918 | Co | 2013 | | 11,512,328 | 126 | 2 | 38,551 | 4,496.2 | 11,416 | 134,390 ( 9.7) | 1 | 1,084,984 | 1,084,984 | 29.24 | 907 | 353 | 34 | 6 | 1 |

**Table S2.** Virulence and VSP genes. Gene count in Prokka annotation. The presence of virulence genes is based on the `Prokka` annotation without reference where the *vsp* gene count is derived from the reference-based annotation.

Virulence Gene Information

| Bio Information | infC | tuf | fusA | rpsL | rpsG | rpmI | rplT | rpoB | rpoC | gyrA | gyrB | ulaB | ulaC | oppD | uvrC | pta | secA | tkt | xerC | eno2 | ldh1 | gls(C) | malF | pyk | ndh | clpC |
|---|---|---|---|---|---|---|---|---|---|---|---|---|---|---|---|---|---|---|---|---|---|---|---|---|---|---|
| Ref 8790 | 1 | 1 | 1 | 1 | 1 | 1 | 1 | 0 | 1 | 1 | 0 | 0 | 0 | 0 | 0 | 2 | 1 | 0 | 0 | 0 | 1 | 0 | 0 | 1 | 0 | 0 |
| Ref PG45 | 1 | 1 | 1 | 1 | 1 | 1 | 1 | 1 | 1 | 1 | 1 | 0 | 0 | 1 | 1 | 2 | 1 | 1 | 0 | 0 | 1 | 0 | 0 | 1 | 0 | 0 |
| DL422_88 | 1 | 1 | 1 | 1 | 1 | 1 | 1 | 1 | 1 | 1 | 1 | 1 | 1 | 1 | 1 | 2 | 1 | 1 | 2 | 1 | 1 | 0 | 0 | 0 | 0 | 0 |
| 15DD0210 | 1 | 1 | 1 | 1 | 1 | 1 | 1 | 1 | 1 | 1 | 1 | 1 | 1 | 1 | 2 | 2 | 1 | 1 | 3 | 1 | 1 | 0 | 0 | 0 | 0 | 0 |
| 15DD0261 | 1 | 1 | 1 | 1 | 1 | 1 | 1 | 1 | 1 | 1 | 1 | 1 | 1 | 1 | 1 | 2 | 1 | 1 | 4 | 1 | 1 | 0 | 0 | 0 | 0 | 0 |
| 15DD0240 | 1 | 1 | 1 | 1 | 1 | 1 | 1 | 1 | 1 | 1 | 1 | 1 | 1 | 1 | 1 | 2 | 1 | 1 | 3 | 1 | 1 | 0 | 0 | 0 | 0 | 0 |
| 15DD0263 | 1 | 1 | 1 | 1 | 1 | 1 | 1 | 1 | 1 | 1 | 2 | 1 | 1 | 1 | 1 | 2 | 1 | 1 | 4 | 1 | 1 | 0 | 0 | 0 | 0 | 0 |
| 17DD0020 | 2 | 1 | 1 | 1 | 1 | 1 | 1 | 1 | 1 | 1 | 1 | 1 | 1 | 1 | 1 | 2 | 1 | 1 | 3 | 1 | 1 | 0 | 0 | 0 | 0 | 0 |
| DL81_99 | 1 | 1 | 1 | 1 | 1 | 1 | 1 | 1 | 1 | 1 | 1 | 1 | 1 | 1 | 1 | 2 | 1 | 1 | 4 | 1 | 1 | 0 | 0 | 0 | 0 | 0 |
| 17DD0007 | 1 | 1 | 1 | 1 | 1 | 1 | 1 | 1 | 1 | 1 | 1 | 1 | 1 | 1 | 1 | 2 | 1 | 1 | 3 | 1 | 1 | 0 | 0 | 0 | 0 | 0 |
| 16DD0054 | 1 | 1 | 1 | 1 | 1 | 1 | 1 | 1 | 1 | 1 | 1 | 1 | 1 | 1 | 1 | 2 | 1 | 1 | 3 | 1 | 1 | 0 | 0 | 0 | 0 | 0 |
| 15DD0250 | 1 | 1 | 1 | 1 | 1 | 1 | 1 | 1 | 1 | 1 | 1 | 1 | 1 | 1 | 1 | 2 | 1 | 1 | 3 | 1 | 1 | 0 | 0 | 0 | 0 | 0 |
| 15DD0163 | 1 | 1 | 1 | 1 | 1 | 1 | 1 | 1 | 1 | 1 | 1 | 1 | 1 | 1 | 1 | 2 | 1 | 1 | 4 | 1 | 1 | 0 | 0 | 0 | 0 | 0 |
| 15DD0207 | 1 | 1 | 1 | 1 | 1 | 1 | 1 | 1 | 1 | 1 | 1 | 1 | 1 | 1 | 1 | 2 | 1 | 1 | 4 | 1 | 1 | 0 | 0 | 0 | 0 | 0 |
| 16DD0100 | 1 | 1 | 1 | 1 | 2 | 1 | 1 | 1 | 1 | 1 | 1 | 1 | 1 | 1 | 1 | 2 | 1 | 1 | 3 | 1 | 1 | 0 | 0 | 0 | 0 | 0 |
| 15DD0123 | 1 | 1 | 1 | 1 | 1 | 1 | 1 | 1 | 1 | 1 | 1 | 1 | 1 | 1 | 1 | 2 | 1 | 1 | 3 | 1 | 1 | 0 | 0 | 0 | 0 | 0 |
| 15DD0140 | 1 | 1 | 1 | 1 | 1 | 1 | 1 | 1 | 1 | 1 | 1 | 1 | 1 | 1 | 1 | 2 | 1 | 1 | 3 | 1 | 1 | 0 | 0 | 0 | 0 | 0 |
| 15DD0233 | 1 | 1 | 1 | 1 | 1 | 1 | 1 | 1 | 1 | 1 | 1 | 1 | 1 | 1 | 1 | 2 | 1 | 1 | 3 | 1 | 1 | 0 | 0 | 0 | 0 | 0 |
| 16DD0001 | 1 | 1 | 1 | 1 | 1 | 1 | 1 | 1 | 1 | 1 | 1 | 1 | 1 | 1 | 1 | 2 | 1 | 1 | 3 | 1 | 1 | 0 | 0 | 0 | 0 | 0 |
| 15DD0228 | 1 | 1 | 1 | 1 | 1 | 1 | 1 | 1 | 1 | 1 | 1 | 1 | 1 | 1 | 1 | 2 | 1 | 1 | 3 | 1 | 1 | 0 | 0 | 0 | 0 | 0 |
| 15DD0165 | 1 | 1 | 1 | 1 | 1 | 1 | 1 | 1 | 1 | 1 | 1 | 1 | 1 | 1 | 1 | 2 | 1 | 1 | 3 | 1 | 1 | 0 | 0 | 0 | 0 | 0 |
| 15DD0249 | 1 | 1 | 1 | 1 | 1 | 1 | 1 | 1 | 1 | 1 | 1 | 1 | 1 | 1 | 1 | 2 | 1 | 1 | 2 | 1 | 1 | 0 | 0 | 0 | 0 | 0 |
| 11DD0261 | 1 | 1 | 1 | 1 | 1 | 1 | 1 | 3 | 2 | 1 | 1 | 1 | 1 | 1 | 1 | 2 | 1 | 1 | 2 | 1 | 1 | 0 | 0 | 0 | 0 | 0 |
| 14DD0148 | 1 | 1 | 1 | 1 | 1 | 1 | 1 | 1 | 1 | 1 | 1 | 1 | 1 | 1 | 1 | 2 | 1 | 1 | 2 | 1 | 0 | 0 | 0 | 0 | 0 | 0 |
| 15DD0161 | 1 | 1 | 1 | 1 | 1 | 1 | 1 | 1 | 1 | 1 | 1 | 1 | 1 | 1 | 1 | 2 | 1 | 1 | 2 | 1 | 1 | 0 | 0 | 0 | 0 | 0 |
| 14DD0475 | 1 | 1 | 1 | 1 | 1 | 1 | 1 | 1 | 1 | 1 | 1 | 1 | 1 | 1 | 1 | 2 | 1 | 1 | 2 | 1 | 1 | 0 | 0 | 0 | 0 | 0 |
| 14DD0156 | 1 | 1 | 1 | 1 | 1 | 1 | 1 | 1 | 1 | 1 | 1 | 1 | 1 | 1 | 1 | 2 | 1 | 1 | 2 | 1 | 1 | 0 | 0 | 0 | 0 | 0 |
| 15DD0238 | 1 | 1 | 1 | 1 | 1 | 1 | 1 | 1 | 1 | 1 | 1 | 1 | 1 | 1 | 1 | 2 | 1 | 1 | 2 | 1 | 1 | 0 | 0 | 0 | 0 | 0 |
| 15DD0234 | 1 | 1 | 1 | 1 | 1 | 1 | 1 | 1 | 1 | 1 | 1 | 1 | 1 | 1 | 1 | 2 | 1 | 1 | 2 | 1 | 1 | 0 | 0 | 0 | 0 | 0 |
| 15DD0218 | 1 | 1 | 1 | 1 | 1 | 1 | 1 | 1 | 1 | 1 | 1 | 1 | 1 | 1 | 1 | 2 | 1 | 1 | 2 | 1 | 1 | 0 | 0 | 0 | 0 | 0 |
| 15DD0164 | 1 | 1 | 1 | 1 | 1 | 1 | 1 | 1 | 1 | 1 | 1 | 1 | 1 | 1 | 1 | 2 | 1 | 1 | 2 | 1 | 1 | 0 | 0 | 0 | 0 | 0 |
| 15DD0141 | 1 | 1 | 1 | 1 | 1 | 1 | 1 | 1 | 1 | 1 | 1 | 1 | 1 | 1 | 1 | 2 | 1 | 1 | 2 | 1 | 1 | 0 | 0 | 0 | 0 | 0 |
| 15DD0160 | 1 | 1 | 1 | 1 | 1 | 1 | 1 | 1 | 1 | 1 | 1 | 1 | 1 | 1 | 1 | 2 | 1 | 1 | 2 | 1 | 1 | 0 | 0 | 0 | 0 | 0 |
| 16DL0615 | 1 | 1 | 1 | 1 | 1 | 1 | 1 | 1 | 1 | 1 | 1 | 1 | 1 | 1 | 1 | 2 | 1 | 1 | 2 | 1 | 1 | 0 | 0 | 0 | 0 | 0 |
| 14DD0147 | 1 | 1 | 1 | 1 | 1 | 1 | 1 | 1 | 1 | 1 | 1 | 1 | 1 | 1 | 1 | 2 | 1 | 1 | 2 | 1 | 1 | 0 | 0 | 0 | 0 | 0 |
| 16DD0186 | 1 | 1 | 1 | 1 | 1 | 1 | 1 | 1 | 1 | 1 | 1 | 1 | 1 | 1 | 1 | 2 | 1 | 1 | 2 | 1 | 1 | 0 | 0 | 0 | 0 | 0 |
| 15DL0124 | 1 | 1 | 1 | 1 | 1 | 1 | 1 | 1 | 1 | 1 | 1 | 1 | 1 | 1 | 1 | 2 | 1 | 1 | 2 | 1 | 1 | 0 | 0 | 0 | 0 | 0 |
| 13DD0918 | 1 | 1 | 1 | 1 | 1 | 1 | 1 | 1 | 1 | 1 | 1 | 1 | 1 | 1 | 1 | 2 | 1 | 1 | 2 | 1 | 1 | 0 | 0 | 0 | 0 | 0 |

VSP 8790-based Annotation

| Bio Information | WP_233435476 | WP_211235702 | WP_136832249 | WP_233435477 | WP_136832641 | WP_136832072 | WP_136832633 | WP_136831985 | WP_136832637 | WP_136832228 | WP_136831481 | WP_136831846 | WP_233435459 | WP_136831431 | WP_136831844 | WP_136832078 | WP_13683208 | WP_136832058 |
|---|---|---|---|---|---|---|---|---|---|---|---|---|---|---|---|---|---|---|
| Ref 8790 | 1 | 1 | 1 | 1 | 1 | 1 | 1 | 1 | 1 | 1 | 1 | 1 | 1 | 1 | 1 | 1 | 1 | 1 |
| Ref PG45 | 0 | 0 | 0 | 0 | 0 | 0 | 0 | 0 | 0 | 0 | 0 | 0 | 0 | 0 | 0 | 0 | 0 | 0 |
| DL422_88 | 1 | 0 | 0 | 0 | 0 | 0 | 0 | 0 | 0 | 1 | 0 | 2 | 1 | 2 | 2 | 1 | 1 | 2 |
| 15DD0210 | 1 | 0 | 0 | 0 | 0 | 0 | 0 | 1 | 0 | 1 | 0 | 0 | 0 | 1 | 1 | 1 | 1 | 2 |
| 15DD0261 | 1 | 1 | 0 | 0 | 0 | 0 | 0 | 1 | 1 | 1 | 0 | 0 | 0 | 1 | 1 | 1 | 1 | 2 |
| 15DD0240 | 1 | 0 | 0 | 0 | 0 | 0 | 1 | 1 | 0 | 1 | 0 | 0 | 0 | 1 | 2 | 1 | 1 | 2 |
| 15DD0263 | 2 | 0 | 0 | 2 | 0 | 0 | 0 | 1 | 0 | 1 | 0 | 0 | 0 | 1 | 1 | 1 | 1 | 2 |
| 17DD0020 | 1 | 0 | 0 | 1 | 0 | 0 | 0 | 0 | 0 | 1 | 0 | 0 | 0 | 1 | 1 | 1 | 1 | 2 |
| DL81_99 | 1 | 1 | 0 | 0 | 0 | 0 | 0 | 1 | 0 | 1 | 0 | 0 | 0 | 1 | 1 | 1 | 1 | 3 |
| 17DD0007 | 1 | 0 | 0 | 0 | 0 | 0 | 0 | 1 | 0 | 1 | 0 | 0 | 0 | 1 | 1 | 1 | 1 | 2 |
| 16DD0054 | 1 | 0 | 0 | 0 | 0 | 0 | 0 | 0 | 0 | 0 | 0 | 0 | 0 | 2 | 1 | 1 | 1 | 2 |
| 15DD0250 | 1 | 0 | 0 | 0 | 0 | 0 | 0 | 0 | 0 | 0 | 0 | 1 | 0 | 2 | 1 | 1 | 1 | 2 |
| 15DD0163 | 2 | 0 | 0 | 1 | 1 | 0 | 0 | 0 | 0 | 0 | 0 | 1 | 0 | 2 | 1 | 1 | 1 | 2 |
| 15DD0207 | 2 | 0 | 0 | 0 | 0 | 0 | 0 | 0 | 0 | 0 | 0 | 1 | 0 | 2 | 1 | 1 | 1 | 2 |
| 16DD0100 | 1 | 0 | 0 | 0 | 0 | 0 | 0 | 0 | 0 | 0 | 0 | 1 | 0 | 2 | 1 | 1 | 1 | 2 |
| 15DD0123 | 1 | 0 | 0 | 0 | 0 | 0 | 0 | 0 | 0 | 0 | 0 | 1 | 0 | 2 | 1 | 1 | 1 | 2 |
| 15DD0140 | 1 | 0 | 0 | 2 | 0 | 0 | 0 | 0 | 0 | 0 | 0 | 1 | 0 | 2 | 1 | 1 | 1 | 3 |
| 15DD0233 | 1 | 0 | 0 | 0 | 0 | 0 | 0 | 0 | 0 | 0 | 0 | 1 | 0 | 2 | 1 | 1 | 1 | 2 |
| 16DD0001 | 1 | 0 | 0 | 1 | 0 | 0 | 0 | 0 | 0 | 0 | 0 | 1 | 0 | 2 | 1 | 2 | 1 | 2 |
| 15DD0228 | 1 | 0 | 0 | 0 | 0 | 0 | 0 | 0 | 0 | 0 | 0 | 1 | 0 | 2 | 1 | 1 | 1 | 2 |
| 15DD0165 | 1 | 0 | 0 | 0 | 0 | 0 | 0 | 0 | 0 | 0 | 0 | 1 | 0 | 2 | 1 | 1 | 1 | 2 |
| 15DD0249 | 1 | 0 | 0 | 0 | 0 | 0 | 0 | 1 | 1 | 0 | 0 | 0 | 0 | 2 | 0 | 1 | 1 | 2 |
| 11DD0261 | 1 | 0 | 0 | 0 | 0 | 0 | 0 | 1 | 1 | 0 | 0 | 1 | 0 | 2 | 1 | 1 | 1 | 2 |
| 14DD0148 | 1 | 0 | 0 | 0 | 0 | 0 | 1 | 1 | 1 | 0 | 0 | 1 | 0 | 2 | 1 | 1 | 1 | 2 |
| 15DD0161 | 1 | 0 | 0 | 0 | 0 | 0 | 1 | 0 | 1 | 0 | 0 | 1 | 0 | 2 | 1 | 1 | 1 | 2 |
| 14DD0475 | 1 | 0 | 0 | 0 | 0 | 0 | 0 | 1 | 1 | 0 | 0 | 1 | 0 | 2 | 1 | 1 | 1 | 2 |
| 14DD0156 | 1 | 0 | 0 | 0 | 0 | 0 | 0 | 1 | 1 | 0 | 0 | 1 | 0 | 2 | 1 | 1 | 1 | 2 |
| 15DD0238 | 1 | 0 | 0 | 0 | 0 | 0 | 0 | 1 | 1 | 0 | 0 | 1 | 0 | 2 | 1 | 1 | 1 | 2 |
| 15DD0234 | 1 | 0 | 0 | 0 | 0 | 0 | 0 | 1 | 1 | 0 | 0 | 2 | 0 | 2 | 1 | 1 | 1 | 2 |
| 15DD0218 | 1 | 0 | 0 | 0 | 0 | 0 | 0 | 0 | 1 | 0 | 0 | 1 | 0 | 2 | 1 | 1 | 1 | 2 |
| 15DD0164 | 1 | 0 | 0 | 0 | 0 | 0 | 0 | 0 | 2 | 0 | 0 | 1 | 0 | 2 | 1 | 1 | 1 | 2 |
| 15DD0141 | 1 | 0 | 0 | 0 | 0 | 0 | 0 | 1 | 0 | 0 | 0 | 1 | 0 | 2 | 1 | 1 | 1 | 2 |
| 15DD0160 | 1 | 0 | 0 | 0 | 0 | 0 | 0 | 0 | 1 | 0 | 0 | 0 | 0 | 2 | 1 | 1 | 1 | 2 |
| 16DL0615 | 1 | 0 | 0 | 0 | 0 | 0 | 0 | 1 | 1 | 0 | 0 | 1 | 0 | 2 | 2 | 2 | 1 | 2 |
| 14DD0147 | 0 | 0 | 0 | 0 | 0 | 0 | 0 | 1 | 1 | 0 | 0 | 1 | 0 | 2 | 1 | 1 | 1 | 2 |
| 16DD0186 | 1 | 0 | 0 | 0 | 0 | 0 | 0 | 1 | 1 | 0 | 0 | 1 | 0 | 2 | 1 | 1 | 1 | 2 |
| 15DL0124 | 1 | 0 | 0 | 0 | 0 | 0 | 0 | 1 | 1 | 0 | 0 | 1 | 0 | 2 | 1 | 1 | 1 | 2 |
| 13DD0918 | 1 | 0 | 0 | 0 | 0 | 0 | 0 | 1 | 1 | 0 | 0 | 1 | 0 | 2 | 1 | 1 | 1 | 2 |

VSP PG45-based Annotation

| Bio Information | A | B | E | F | G | H | I | J | K | L | M | N | O |
|---|---|---|---|---|---|---|---|---|---|---|---|---|---|
| Ref 8790 | 0 | 0 | 0 | 0 | 0 | 0 | 0 | 0 | 0 | 0 | 0 | 0 | 0 |
| Ref PG45 | 1 | 1 | 1 | 1 | 1 | 1 | 1 | 1 | 1 | 1 | 1 | 1 | 1 |
| DL422_88 | 1 | 0 | 0 | 0 | 1 | 0 | 0 | 0 | 0 | 1 | 0 | 0 | 0 |
| 15DD0210 | 2 | 2 | 0 | 0 | 1 | 1 | 0 | 0 | 1 | 0 | 0 | 0 | 0 |
| 15DD0261 | 0 | 0 | 0 | 0 | 2 | 0 | 0 | 1 | 1 | 1 | 0 | 0 | 1 |
| 15DD0240 | 1 | 0 | 0 | 0 | 1 | 1 | 1 | 0 | 2 | 0 | 0 | 1 | 0 |
| 15DD0263 | 9 | 0 | 0 | 0 | 1 | 0 | 0 | 0 | 1 | 3 | 0 | 0 | 0 |
| 17DD0020 | 4 | 0 | 0 | 0 | 0 | 0 | 0 | 0 | 0 | 1 | 0 | 0 | 0 |
| DL81_99 | 7 | 0 | 0 | 0 | 0 | 0 | 0 | 0 | 1 | 1 | 0 | 0 | 0 |
| 17DD0007 | 5 | 0 | 0 | 0 | 1 | 1 | 0 | 0 | 1 | 1 | 0 | 0 | 0 |
| 16DD0054 | 0 | 0 | 0 | 0 | 2 | 0 | 2 | 1 | 0 | 0 | 0 | 2 | 2 |
| 15DD0250 | 0 | 0 | 0 | 0 | 2 | 1 | 1 | 1 | 1 | 1 | 0 | 2 | 1 |
| 15DD0163 | 1 | 0 | 0 | 0 | 1 | 4 | 0 | 1 | 3 | 0 | 0 | 4 | 2 |
| 15DD0207 | 0 | 0 | 0 | 0 | 0 | 4 | 0 | 1 | 4 | 0 | 0 | 4 | 3 |
| 16DD0100 | 0 | 0 | 0 | 0 | 1 | 1 | 0 | 1 | 1 | 1 | 0 | 3 | 2 |
| 15DD0123 | 0 | 0 | 0 | 0 | 2 | 1 | 1 | 1 | 1 | 1 | 0 | 2 | 2 |
| 15DD0140 | 0 | 0 | 0 | 0 | 2 | 1 | 1 | 2 | 1 | 1 | 0 | 2 | 3 |
| 15DD0233 | 0 | 0 | 0 | 0 | 2 | 1 | 1 | 1 | 1 | 1 | 0 | 2 | 2 |
| 16DD0001 | 0 | 0 | 0 | 0 | 2 | 2 | 1 | 1 | 1 | 1 | 0 | 2 | 3 |
| 15DD0228 | 1 | 0 | 0 | 0 | 1 | 0 | 0 | 1 | 0 | 1 | 0 | 1 | 0 |
| 15DD0165 | 0 | 0 | 0 | 0 | 2 | 1 | 1 | 1 | 1 | 1 | 0 | 2 | 3 |
| 15DD0249 | 1 | 0 | 1 | 0 | 2 | 0 | 1 | 0 | 2 | 1 | 1 | 1 | 0 |
| 11DD0261 | 1 | 0 | 1 | 0 | 2 | 0 | 1 | 0 | 1 | 1 | 1 | 1 | 0 |
| 14DD0148 | 3 | 0 | 2 | 0 | 2 | 0 | 1 | 0 | 1 | 1 | 1 | 4 | 0 |
| 15DD0161 | 1 | 0 | 1 | 0 | 1 | 1 | 1 | 0 | 1 | 1 | 1 | 1 | 0 |
| 14DD0475 | 1 | 0 | 1 | 0 | 3 | 1 | 1 | 0 | 1 | 1 | 1 | 1 | 0 |
| 14DD0156 | 0 | 0 | 0 | 0 | 1 | 0 | 1 | 0 | 1 | 1 | 0 | 1 | 0 |
| 15DD0238 | 1 | 0 | 1 | 0 | 1 | 0 | 1 | 0 | 1 | 1 | 1 | 1 | 1 |
| 15DD0234 | 1 | 0 | 1 | 0 | 2 | 0 | 1 | 0 | 1 | 1 | 1 | 0 | 0 |
| 15DD0218 | 2 | 0 | 0 | 0 | 3 | 0 | 0 | 0 | 2 | 2 | 1 | 2 | 0 |
| 15DD0164 | 1 | 0 | 0 | 0 | 3 | 0 | 0 | 0 | 3 | 2 | 2 | 3 | 0 |
| 15DD0141 | 0 | 0 | 0 | 0 | 1 | 1 | 1 | 0 | 1 | 1 | 0 | 0 | 0 |
| 15DD0160 | 1 | 0 | 0 | 0 | 2 | 0 | 0 | 0 | 1 | 1 | 1 | 1 | 0 |
| 16DL0615 | 1 | 0 | 1 | 0 | 2 | 0 | 1 | 0 | 2 | 1 | 1 | 1 | 0 |
| 14DD0147 | 1 | 0 | 0 | 0 | 0 | 0 | 1 | 0 | 1 | 1 | 0 | 1 | 0 |
| 16DD0186 | 1 | 0 | 1 | 0 | 2 | 0 | 1 | 0 | 0 | 1 | 0 | 1 | 0 |
| 15DL0124 | 1 | 0 | 1 | 0 | 1 | 0 | 1 | 0 | 1 | 1 | 0 | 2 | 0 |
| 13DD0918 | 2 | 0 | 1 | 0 | 3 | 0 | 1 | 0 | 2 | 2 | 1 | 1 | 0 |

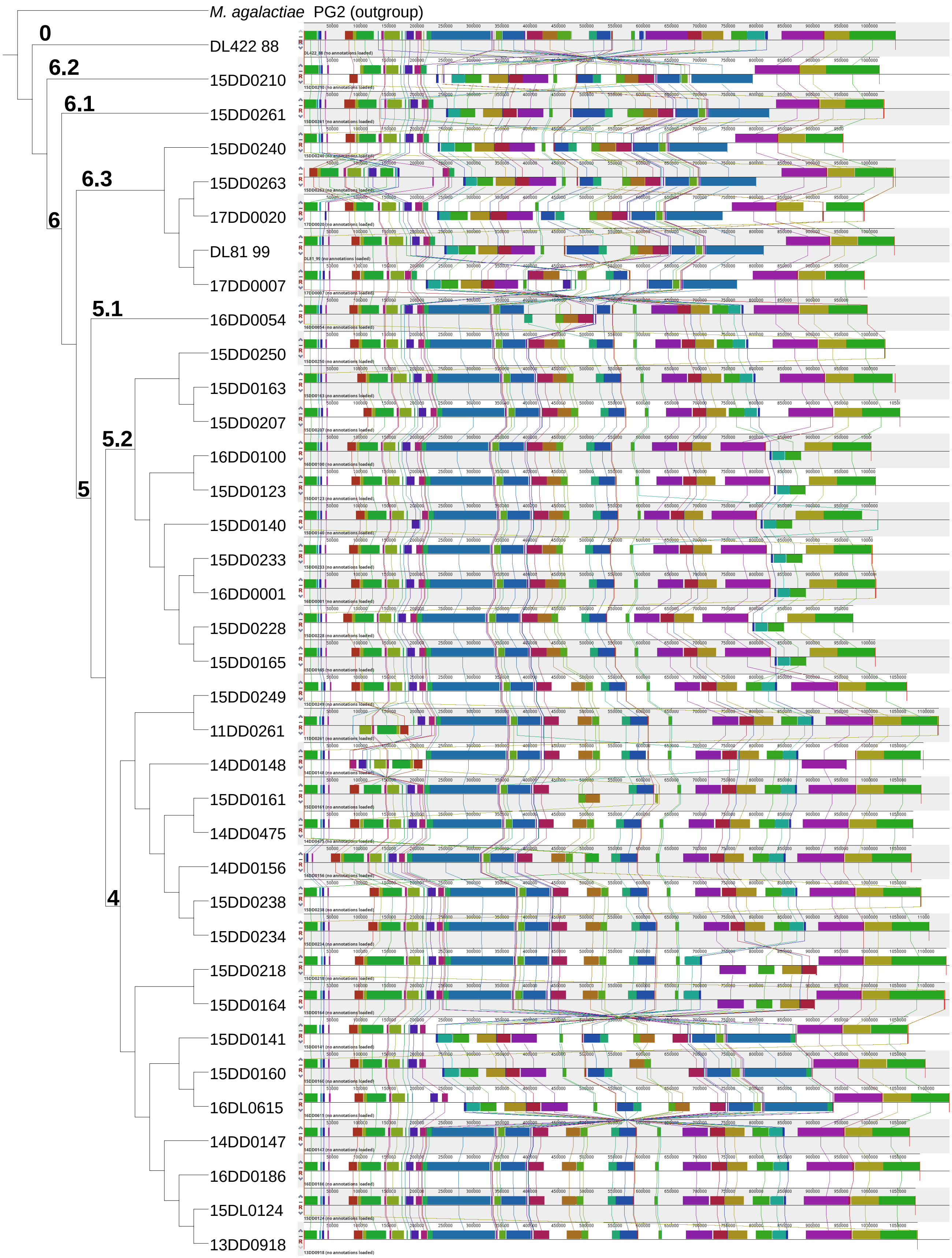


**Figure S2.** Recombinations within our 36 assemblies. Constructing whole-genome alignments with `Mauve` revealed the same organization in the 5' and 3' regions, but several changes in between, consistent with phylogeny. We observed rearrangements conserved in subtrees, indicating a true rearrangement and not an assembly error. The alignment of representative assemblies is shown in Fig. 4.

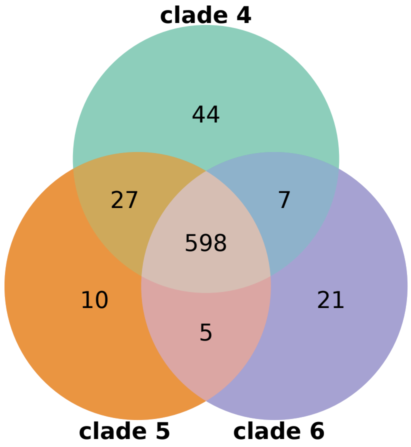


**Figure S3.** Schematic visualisation of the pairwise gene association analysis. Pairwise gene association analysis detected 114 significant genes in total; 49 genes linked to clade 4 (44 positive and 5 negative), 17 to clade 5 (10 positive and 7 negative), and 48 to clade 6 (21 positive and 27 negative).
